# Full-length structure and heme binding in the transcriptional regulator HcpR

**DOI:** 10.1101/2024.09.06.611725

**Authors:** B. Ross Belvin, Faik N. Musayev, Carlos Escalante, Janina P. Lewis

## Abstract

HcpR is a CRP-family transcriptional regulator found in many Gram-negative anaerobic bacteria. In the perio-pathogen *Porphyromonas gingivalis,* HcpR is crucial for the response to reactive nitrogen species such as nitric oxide (NO). Binding of NO to the heme group of HcpR leads to transcription of the redox enzyme Hcp. However, the molecular mechanisms of heme binding to HcpR remain unknown. In this study we present the 2.3Å structure of the *P. gingivalis* HcpR. Interdomain interactions present in the structure help to form a hydrophobic pocket in the N-terminal sensing domain. A comparison analysis with other CRP-family members reveals that the molecular mechanisms of HcpR-mediated regulation may be distinct from other family members. Using docking studies, we identify a putative heme binding site in the sensing domain. *In vitro* complementation and mutagenesis studies verify Met68 as an important residue in activation of HcpR. Finally, heme binding studies with purified forms of recombinant HcpR support Met68 and His149 residues as important for proper heme coordination in HcpR.

## Introduction

HcpR is a sensor-transcriptional regulator of the nitrosative stress response. Found in many Gram-negative anaerobic bacteria, it regulates the expression of *hcp*, an enzyme crucial for survival in response to nitric oxide (NO) and other forms of nitrosative stress ^1,2^. HcpR belongs to the cAMP receptor (CRP) family of proteins of which the catabolite gene activator protein (CAP) and the CO sensing protein CooA are also members ^3,4^. This large family of proteins sense changes in the intra-cellular environment by binding effector molecules that allosterically activate the protein for DNA binding ^5^. The CRP family contains a significant number of sub-families that have evolved to sense a wide variety of effector molecules and utilize different cofactors ^5^. One of these sub-families, the DNR-HcpR group has evolved to sense NO using metal cofactors such as heme and is phylogenetically distinct from other heme containing CAP family members such as CooA ^6^. Although not all HcpRs have been demonstrated to bind heme, a growing subset have been characterized as heme binding proteins ^7^.

HcpR is crucial for the survival of the periodontal pathogen *Porphyromonas gingivalis* ^8^. In the periodontal pocket, *P. gingivalis* is exposed to numerous nitrosative stresses, including immune cell derived NO as well as elevated levels of nitrite from saliva ^9,10^. Loss of *hcpR* or *hcp* sensitizes the bacteria to physiological concentrations of nitrite and NO and causes decreased survival with host cells and *in vivo* using mouse models ^8,11–13^. Previously we have demonstrated that HcpR from *P. gingivalis* can sense nanomolar concentrations of NO and upregulates the expression of *hcp* in response to these stresses ^13^. Furthermore, we have shown that binding of HcpR to the *hcp* promoter is dependent on heme ^8^.

In heme-based sensor proteins, the heme iron complex serves as the binding site for gaseous molecules such as NO, CO, or O_2_. Displacement of an axial side chain that coordinates the heme iron is often utilized for allosteric activation of heme proteins ^14,15^. In HcpR, the heme exists in a 6-coordinate system in the unbound when no gas is bound; upon addition of NO, this switches to a 5-coordinate system ^13^. The capability of NO to displace the proximal side chain is a distinctive facet of the NO-heme interaction that is not typically seen in CO or O_2_ heme sensor proteins ^15^. The 5-coordinate NO bound heme in NO sensor proteins such as HcpR and DNR distinguishes it from the CO sensor CooA, which is a 6-coordinate system in the CO bound state ^16,17^. The different nature of the gas binding properties has implications for the mechanism of NO dependent activation in HcpR.

So far, the identity of the axial side chains in HcpR is not known. Furthermore, the nature of heme binding to HcpR (the location of the exact heme binding pocket) is not understood. Previously we have solved the structure of the sensing domain of HcpR (ΔCHcpR), that lacks the (highly flexible) C-terminal DNA binding domain ^8^. While the truncated structure confirms the inclusion of HcpR as CRP-family regulator, information on inter-domain interactions and how this may influence the heme binding nature of the protein was lost. In this study, we present the structure of the full-length HcpR (FL HcpR). Using a docking approach supported by site-directed mutagenesis and heme binding studies, we identify a putative heme-binding site and important residues in heme coordination. A comparison of HcpR with the structures of CooA and CAP reveals a novel mechanism that may control the NO-mediated activation of HcpR.

## Materials and Methods

### Generation of recombinant proteins

The full length of the HcpR-encoding gene (PG1053) was cloned into the pET21d vector at the *Bam*HI and *Xho*I sites using gene specific primers with complimentary restriction site overhangs. Vector and PCR fragments were cleaved, purified, ligated and then transformed into DH5α *E. coli* for screening. Positive clones were identified and whole plasmid sequencing was subsequently performed to confirm the proper genetic content. The recombinant *hcpR-*pET21 vector was transformed into BL21(DE3) or Lemo21(DE3) cells (New England Biolabs) to enable inducible protein expression.

For recombinant HcpR expression, overnight culture was used to inoculate 1 L of LB broth. Cultures were grown at 37°C with antibiotics (50 ug/mL carbenicillin) and shaking until cultures reached mid-log phase (0.5-0.7 OD600) and expression of HcpR was induced using 1mM IPTG overnight at 37°C. HcpR was affinity purified using Ni-NTA agarose (Qiagen) or TALON Cobalt resin (ThermoFisher). Briefly, cells were lysed using CellLytic B reagent (Millipore-Sigma) and cell supernatant was added to column containing metal affinity resin. Columns were washed with a 50mM Na_2_H_1_PO_4_, 300mM NaCl, 20mM imidazole, pH 8.0 buffer. For elution, imidazole concentration was increased to 250mM. For crystallization, HcpR was further purified via gel exclusion chromatography on a Superdex 75 column. Purification of recombinant mutant HcpRs was performed identically to the wildtype HcpR.

For heme reconstitution, purified HcpR was concentrated and desalted into a 25mM Phosphate, 150mM NaCl, 1mM TCEP pH 7.4 buffer using a PD-10 desalting column. A 1.1 molar excess of heme was added to purified HcpR and allowed to incubate overnight. To remove unbound heme, HcpR was dialyzed overnight and/or desalted using a spin column.

### Crystallization and structure determination

Freshly prepared HcpR (15.1mg/ml) in 25mM Tris-HCl, 150mM NaCl, 1mM TCEP, pH 7.0 was used to screen for initial crystallization condition using a wide range of commercially available screening kits with the Gryphon crystallization robot (Art Robbins) employing the sitting-drop vapor-diffusion method at 20°C. 0.3μl of protein solution and 0.3 μl of reservoir were mixed to equilibrate against 60μl reservoir solution. Crystals appeared in one week under several conditions (K/Na tartrate, Na formate, Na citrate tribasic and PEG-3350) at pH range 4.6-6.5. We have crystallized high diffraction grade single crystal of wild type HcpR at low pH (4.6). Crystallization condition was 0.1M sodium acetate, pH 4.6 and 1.8M sodium formate. The crystals diffracted up to 2.3Å resolution and belonged to space group P212121 with the unit cell dimensions of a = 106.260, b = 106.303, and c = 116.773Å. Crystals were fished from the drops and cryo-protected by mother liquor varied concentration of glycerol (10-20%). Subsequently, fished crystals were flash-cooled in a cryogenic nitrogen stream. X-ray diffraction data were collected at 100K using Rigaku MicroMax-007 HF X-ray generator and an EIGER R 4M detector. The data was processed with *CrysAlis^Pro^* 40.64.69a (Rigaku) and the CCP4 suite of programs.

### Model building and refinement

The crystal structure was solved with *Phaser* in the *Phenix* software package using the sensing domain of HcpR (PDB entry: 6NP6) as the search model. Auto model building resulted in an initial model of 572 amino-acid residues built with R_work_ 28.21% and R_free_ 31.71%. The structure was refined with the Phenix software along with manual model building using the graphics program Coot. After several cycles of refinement and manual model building positive electron density belonging to the C-terminal domain became visible. It was possible to fit helix-turn-helix motif of DNA binding domain. The structure was refined to 2.3Å resolution with a final R_work_ of 21.57% and R_free_ of 26.47%.

### Docking studies

Structural models of the heme group and the FL-HcpR dimer were prepared for docking calculations using the Graphical User Interface AutoDock Tools (ADT). The grid size was set to 40 x 40 x 40 xyz points with grid spacings of 0.375 Å and grid center was placed at x= 111.826, y= 42.032, z= 151.488. Other parameters were maintained at the default configuration. The two pqt files were imported into AutoDock Vina tool based on a fixed geometry of HcpR and Heme. The result with the lowest energy of binding (−8.3 kcal/mol) was selected for further analysis in ChimeraX.

### UV-Vis spectrum studies

The spectrum of heme-reconstituted HcpR was recorded using a Genesys 150 UV-Vis spectrophotometer (ThermoFisher) in a gas tight 1 cm quartz cuvette in 25mM sodium phosphate, 150mM NaCl, 1mM TCEP, pH 7.4 buffer. The ferrous form of heme was obtained by adding an excess of sodium dithionite. To obtain anaerobic samples, purified HcpR was incubated in an anaerobic chamber overnight.

### Bacterial strains and growth conditions

*P. gingivalis* W83 (strain V2802) was grown and maintained in an atmosphere consisting of 80% N2, 10% CO2, and 10% H2 in an anaerobic chamber (Coy Laboratories). The *hcpR* deficient *P. gingivalis* (V2807) was derived previously ^8^. *P. gingivalis* strains were grown on TSA-blood agar plates (Tryptic Soy Agar, 5% Sheep blood) or in tryptic soy broth (TSB) supplemented with hemin (5 ug/mL) and Vitamin K (1 ug/mL). Plasmid containing strains were maintained on plates or grown in TSB broth containing tetracycline (0.5 ug/mL).

### Cloning and site directed mutagenesis of *P. gingivalis hcpR* complement plasmids

The *P. gingivalis hcpR* deficient strain (V2807) was complemented using the pG108 plasmid ^18^. A copy of the *hcpR* gene was synthesized downstream of the *ermF* promoter to create a construct that will constitutively express HcpR. The construct was cloned into the pG108 vector at the *Sph*I and *Bam*HI restriction sites to create the pG108-*hcpR* plasmid. Single amino acid mutants were created using the QuikChange II XL site directed mutagenesis kit (Agilent). Primers were designed to introduce mutations at Met68, His149 and L156 to convert them to Ala or a stop-codon. Primers used for this study are listed in supplemental table. Clones generated from mutagenesis were submitted for full plasmid sequencing and positive clones were used to complement the *hcpR* deficient V2807 strain.

### Electroporation and complementation of HcpR-deficient *P. gingivalis*

Fresh bacterial cultures on blood plates were used to inoculate 5 ml of BHI broth culture overnight at 37°C. The next day, this culture was diluted 1:10 with fresh media and grown to mid-log phase (0.5-0.6 OD_600_). Cells were harvested by centrifugation at 8000x g for 15 minutes at 4°C, washed in 25mL cold electroporation buffer (10% glycerol, 1mM MgCl2) and suspended in ∼250 µL EP buffer. To a pre-chilled 0.2 cm electroporation cuvette, 50 µL of washed cells and 1-2 ug of pG108-*hcpR* plasmid DNA was added. The cuvette was placed in an electroporation chamber and pulsed using settings at 2.5kV, 5msec, 400 Ω on a BioRad Gene Pulser II. Immediately after electroporation, 500 µL of pre-warmed media was added and cells were grown overnight in the anaerobic chamber. The next day, the bacteria were plated on TSA-blood agar plates supplemented with 0.5ug/mL tetracycline. Colonies that appeared after 5-6 days of incubation were replated and screened for proper genetic content.

### Nitrite growth studies

Overnight cultures of plasmid complemented Δ*hcpR P. gingivalis* started from TSA-blood plates were diluted to on OD_600_ of 0.05. These cultures were supplemented with 0 mM, 1mM, or 2mM sodium nitrite and grown overnight anaerobically at 37°C. The next day the OD6_600_ was measured to assess growth of the *P. gingivalis*strains.

### qRT-PCR

To test HcpR activity *in vitro, P. gingivalis* V2807 complemented with plasmids containing recombinant *hcpR* were grown to mid-log phase in TSB broth and then exposed to 0.2mM nitrite for 15 minutes. The bacterial cells were harvested, RNA was purified using Zymo Quick-RNA mini-prep kit, treated with DNA-free DNAse kit (ThermoFisher) and cDNA was generated using High-Capacity cDNA reverse transcriptase kit (ThermoFisher). Real time qPCR analysis was performed with a SYBR green based detection system on a Quant Studio 3 thermal cycler. Primers used for qPCR: hcp-F: AAAGCTGTCATCGTCCTGCT ; hcp-R: CGATCAGCGTCCGAATATCT ; Pg16s-F: AGGCGGAATTCGTGGTGTAG ; Pg16s-R: TTTGATACCCACGCCTTCGT

### Bioinformatics

Molecular graphics and analyses were performed with UCSF ChimeraX, developed by the Resource for Biocomputing, Visualization, and Informatics at the University of California, San Francisco, with support from National Institutes of Health R01-GM129325 and the Office of Cyber Infrastructure and Computational Biology, National Institute of Allergy and Infectious Diseases ^19,20^. ChimeraX was used to superimpose structures, determine distances, and any potential clashes/contracts. AlphaFold 3 was used to model the DNA binding domains in the active form bound to DNA ^21^. Sequence alignments were constructed using Clustal Omega ^22^. Alignments were portrayed using Jalview ^23^.

## Results

### Overview of the structure of full length HcpR

Crystals of the FL-HcpR belong to the space group P4_1_2_1_2 and diffracted to 2.3 Å. The structure was solved by molecular replacement using the truncated HcpR structure (PDB ID 6NP6). The asymmetric unit contains 2 dimers with chains A, B, C, and D. There was no significant loss of electron density along the backbone of the structure, and all 228 amino acids of the gene CDS are accounted for. In subunits A and C, the uncleaved cloning tail containing polyhistidine and TEV sites can also be observed on the map. The data collection and refinement statistics are summarized in Table 1.

**Table 1:** Data Collection and refinement statistics.

| **Data Collection Statistics** |  |
| --- | --- |
| Space group | P 2<sub>1</sub> 2<sub>1</sub> 2<sub>1</sub> |
| Unit-cell *a*, *b*, *c* (Å) | 106.260, 106.303, 116.773 |
| Resolution (Å) | 29.13–2.30 (2.36 – 2.30) |
| Total reflections | 711884 (55391) |
| Unique reflections | 59441 (4590) |
| Redundancy | 12.0 (12.1) |
| Completeness (%) | 99.9 (100.0) |
| Average *I*/σ(*I*) | 33.7 (4.6) |
| R<sub>merge</sub> (%)<sup>a</sup> | 5.9 (60.1) |
| **Refinement Statistics** |  |
| Resolution (Å) | 29.13-2.30 (2.38-2.30) |
| No. of reflections | 59146 (2542) |
| R<sub>work</sub> (%) | 21.57 (26.87) |
| R<sub>free</sub> (%)<sup>b</sup> | 26.47 (32.05) |
| R.m.s.d. bonds (Å) | 0.002 |
| R.m.s.d. angles (°) | 0.5800 |
| Dihedral angles |  |
| Most favored (%) | 94.4 |
| Allowed (%) | 4.6 |
| **Average B (Å<sup>2</sup>) / atoms** |  |
| All atoms | 69.00 |
| Protein | 71.01 |
| Ligand | 56.86 |
| Solvent | 51.31 |
| **Number of non-hydrogen atoms** | 7632 |
| Macromolecules | 7221 |
| Ligands | 48 |
| Water | 363 |
| Protein residues | 926 |

<sup>a</sup> $R_{\text{merge}} = \frac{\sum_{hkl} \sum_i |I_i(hkl) - \langle I(hkl) \rangle|}{\sum_{hkl} \sum_i I_i(hkl)}$ . <sup>b</sup> $R_{\text{free}}$ was calculated from 5% randomly selected reflection for cross-validation. All other measured reflections were used during refinement.

Structurally, each monomer of HcpR comprises an N-terminal ligand sensing domain (SD) and a C-terminal DNA binding domain (DBD). In agreement with our previous work, HcpR forms a ∼48 kDa homodimer (Fig 1A). In HcpR, the SD encompasses residues M1-L151 (Figure 1A). The DBD extends from residues L156-E228 and forms a winged helix turn helix motif. The dimerization helices (helix E) of each monomer form a leucine zipper motif along residues P127-L151 with interface interactions between symmetry-related Leu, Ile, and Met residues stabilizing the formation of the homodimer (Fig 1B). The leucine zipper and DBD are connected by residues spanning S152-L156, known as the ‘hinge’. This is a metamorphic region that is unfolded in chain A, but forms part of an extended helix E in chain B, spanning residues 127 to 171 (Figure 1C). This hinge property is key to the mechanisms that govern the activation and DNA binding of many CRP family regulators ^24^.

**Figure 1.**
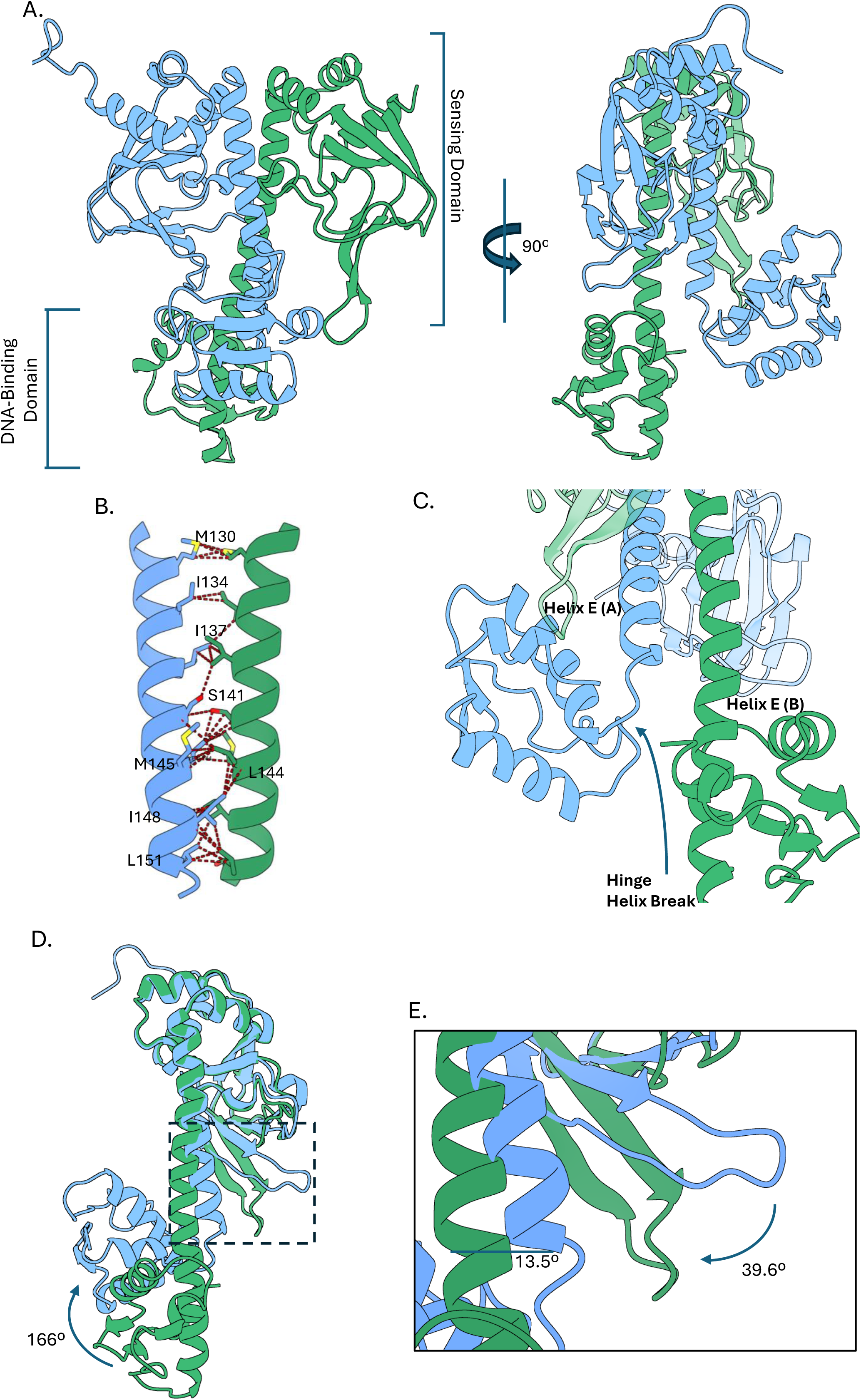
Overview of the crystal structure of full length HcpR. (A) Ribbon diagram of the HcpR shown in forward and 90° side view showing orientation of the N-terminal sensing domain and C-terminal DNA binding domain. Chain A is colored in blue and Chain B is colored in green. The two chains form a homodimer. (B) Dimerization helices of HcpR. The homodimer is stabilized by hydrophobic interactions between symmetry related Met, Leu, and Ile residues. (C) Highlight of the asymmetry of the DNA binding domains of chains A and B. Helix E of chain A is broken at the hinge region at residue 152 where Helix E of chain B is extended through residue 170. (D) Superimposing chains A and B highlights the asymmetry of the DNA binding domains. The DBD of chain A is rotated 166° with respect to chain B. (E) Inset of boxed region in panel (A). The flap region of chain B is angled backwards 39.6 ° create a pocket in the SD of chain B.

The ligand sensing domain of HcpR is composed of 7 β-sheets and 4 α-helices. The β-sheets form a β-barrel or “jelly roll” like fold typically found in the FNR-CRP family of regulators. According to the standard topology of CRP-family regulators, this region consists of 8 β-sheets organized in an anti-parallel fashion. Consistent with what was observed in the crystal structure of the truncated HcpR, the location of the 6^th^ β-sheet is completely distorted, owing to a proline residue (Pro89) creating an extended loop in the β6-β7 turn with a 1 turn helix. The presence of this extended loop region thereby influences the potential size and location of the ligand binding pocket. Pro89 and the region surrounding it is highly conserved in the HcpRs from *Porphyromonas* and *Prevotella* species (Fig. S1), indicating it may play a pivotal role in the formation of the ligand binding pocket.

Superimposing the two HcpR chains identified three structurally dissimilar regions (rmsd 2.14 Å). The DNA binding domain of chain B is rotated by almost 166° with respect to that of chain A (Fig. 1D). One subunit is in an ‘OFF’ state, while the other is in an apparent ‘ON’ state. The second region is in the position of the flap region, with a rotation of 39.6° between them (Fig. 1E). The difference in conformation is generated by the interaction between the β-hairpin of chain A and the DBD of chain B, particularly an interaction between R226 and main chain carbonyl oxygens from V69 and G70 (Fig. S2). The final region is in the second half of the leucine zipper, where they superimpose, forming a 13.5° angle (Fig. 1E).

### Comparison to other CRP-family regulators

A structural comparison with CooA, the only structurally shown heme-binding protein belonging to the CRP family, was carried out to identify the potential heme binding pocket in HcpR (Figure 4). Overall, the secondary structure features and domain architectures are well conserved between the two proteins and they superimpose with a rmsd of 5.45 Å (Fig 2A). In CooA the heme binding pocket is located at the first half of the α-helices forming the leucine zipper. The heme group is coordinated by the N-terminal Pro2 of one subunit and His77 of the second subunit located on the loop connecting β6 and β7 (Fig. 2B). From the superposition, the position of the heme group in HcpR is occupied by the extended loop connecting β6 and β7 which is significantly longer and has a short 1 turn α-helix (Fig. 2B). This extension is also evident in the sequence alignment of HcpR and CooA (Fig. S3). Moreover, in HcpR, no residues are nearby to coordinate with the heme group. Comparison with CAP reveals that the heme and cyclic-AMP binding sites are situated in close but not identical positions (Fig. S4A). In HcpR, the extended loop region once again occupies a similar location, partially occupying the cyclic-AMP binding site in CAP (Fig. S4B). Furthermore, the overall volume of the pocket in CAP is smaller to accommodate the smaller ligand cAMP. Taken together, we conclude that the presence of the long β6-β7 loop suggests that the heme binding pocket in HcpR must be in a different location.

**Figure 2.**
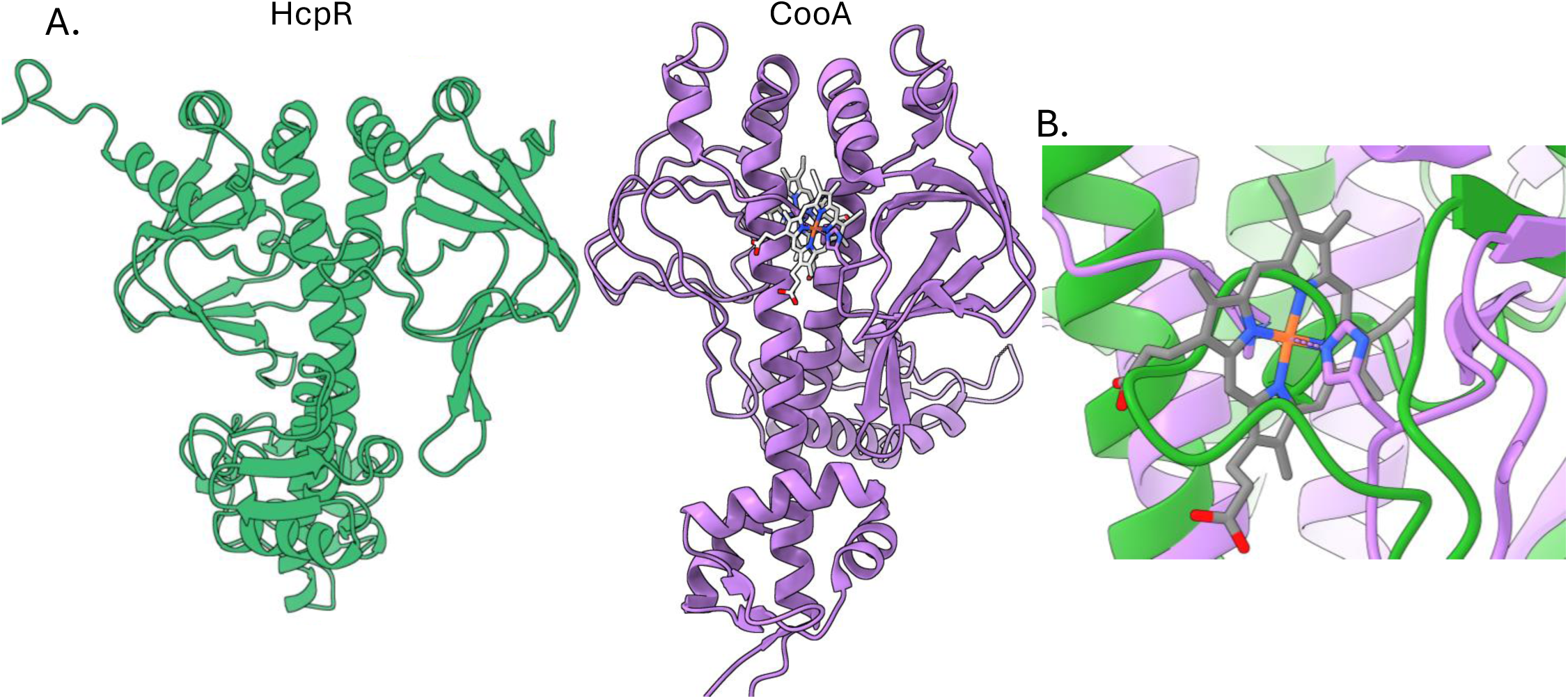
Comparison of the overall fold and conformational differences between HcpR and CooA. (A) Stereo view of HcpR (green) and CooA (purple) dimers depicting the overall organization of each protein. The N-terminal domain of HcpR (extending from residue 1-127) is slightly larger than that of CooA (extending from residues 2-107). (B) Superposition of the HcpR-CooA overlay at the location of CooA binding site. The extended loop region of HcpR (89-93) occupies this region.

**Figure 3.**
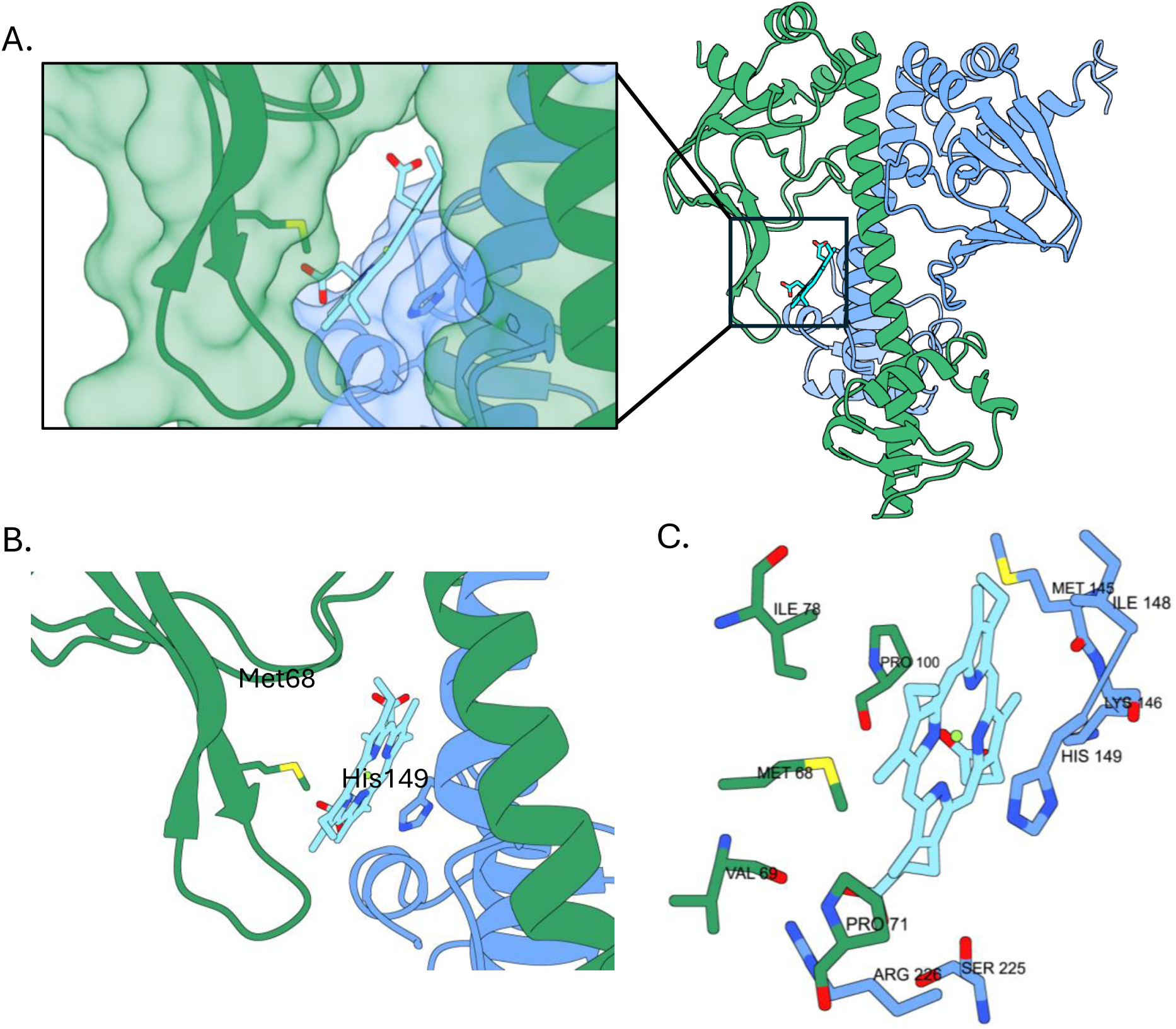
Heme docks into the hydrophobic pocket of HcpR. (A) Overview of heme docking in the HcpR pocket. Heme docks at the interface between chain B SD and middle portion of helix E of chain A. (B) Side chains present in the putative heme pocket. Both Met68 and His149, residues commonly implicated in heme coordination, are optimally located in the pocket to coordinate the heme iron. (C) Side chains present in the heme pocket that form potential contacts with docked heme.

**Figure 4.**
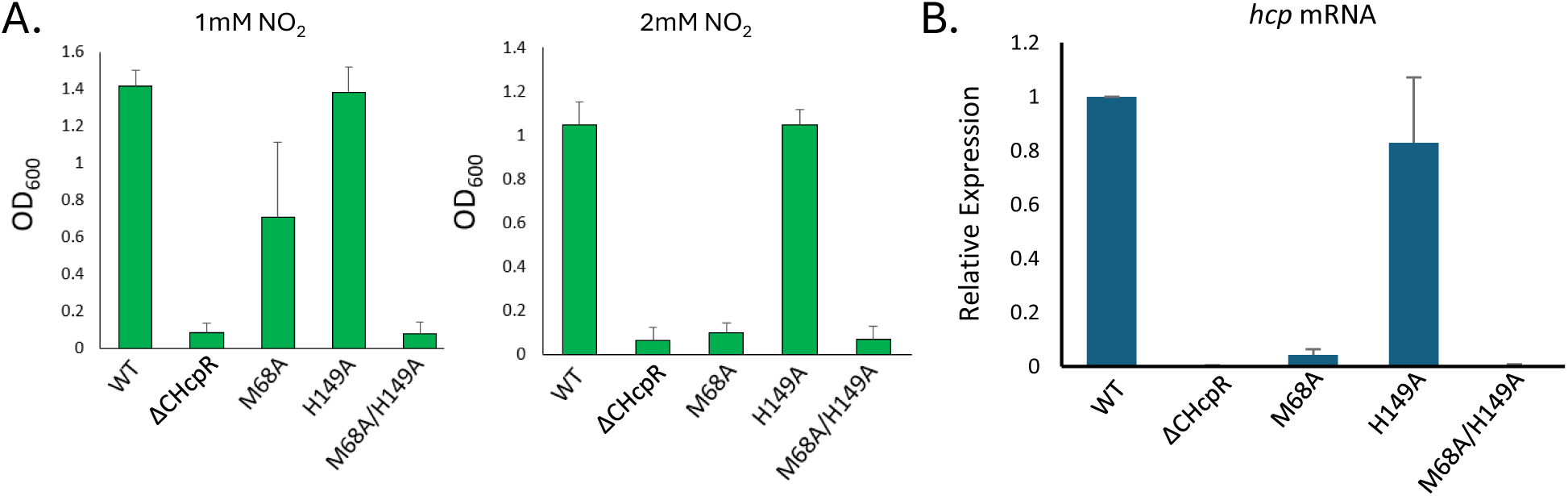
Met68 is important for proper function of HcpR. (A) Growth of ΔHcpR *P. gingivalis* complemented with WT and mutant HcpRs (L156* mutant lacks the DNA binding domain and serves as a negative control). Complemented strains were grown in 1mM NO_2_ or 2mM NO_2_ for 24 hours in Tryptic Soy Broth. After incubation the optical density at 600nm was used to assess growth. (B) Complemented ΔHcpR *P. gingivalis* strains growing in mid-log phase were exposed to 0.2mM NO_2_ for 15 minutes and collected. The level of *hcp* transcript was assessed via qRT-PCR to assess activity of HcpR mutants. Expression levels of *hcp* mRNA are normalized to the 16s subunit mRNA.

### HcpR heme binding pocket

We previously determined the structure of the ΔCHcpR sensing domain and tried to identified heme binding sites. Although our research found two hydrophobic pockets, they were not large enough to accommodate a heme molecule (7). Using resonance Raman spectroscopy, we also determined that heme-bound HcpR is primarily in a 6-coordinate state, with one of the axial bonds being weaker and easily broken. Thus, it is likely that the two side chains involved in coordination are essential for activating HcpR. To search for potential heme-binding sites, we use AutoDock Vina using the structure of the dimer as the target (22). The highest-scoring prediction positioned the heme group at a crevice located at the interface of the two subunits (Fig. 5A). Particularly the DBD of chain A and the flap of the SD of chain B. Crucially, two residues, Met68 and His149, appear primed in the pocket to coordinate the heme iron (Fig. 5B). Apart from the heme iron coordination, many residues from both chains in and around the pocket are in possible positions to form contacts with heme. These include hydrophobic interactions from Val69, Ile78, Pro71, Pro100, Met145, and Ile148, and hydrogen bonding and ionic/salt bridge interactions between the acid groups of heme and Lys146, Ser225, and Arg226 (Fig. 5C). In the crystal structure this pocket is occupied by a portion of the polyhistidine-tail and the TEV protease site that transverses across the tunnel created by the flap of molecule B and the DBD of molecule A (Figure X). The hydrophobic character of the pocket induces the tail interaction.

**Figure 5.**
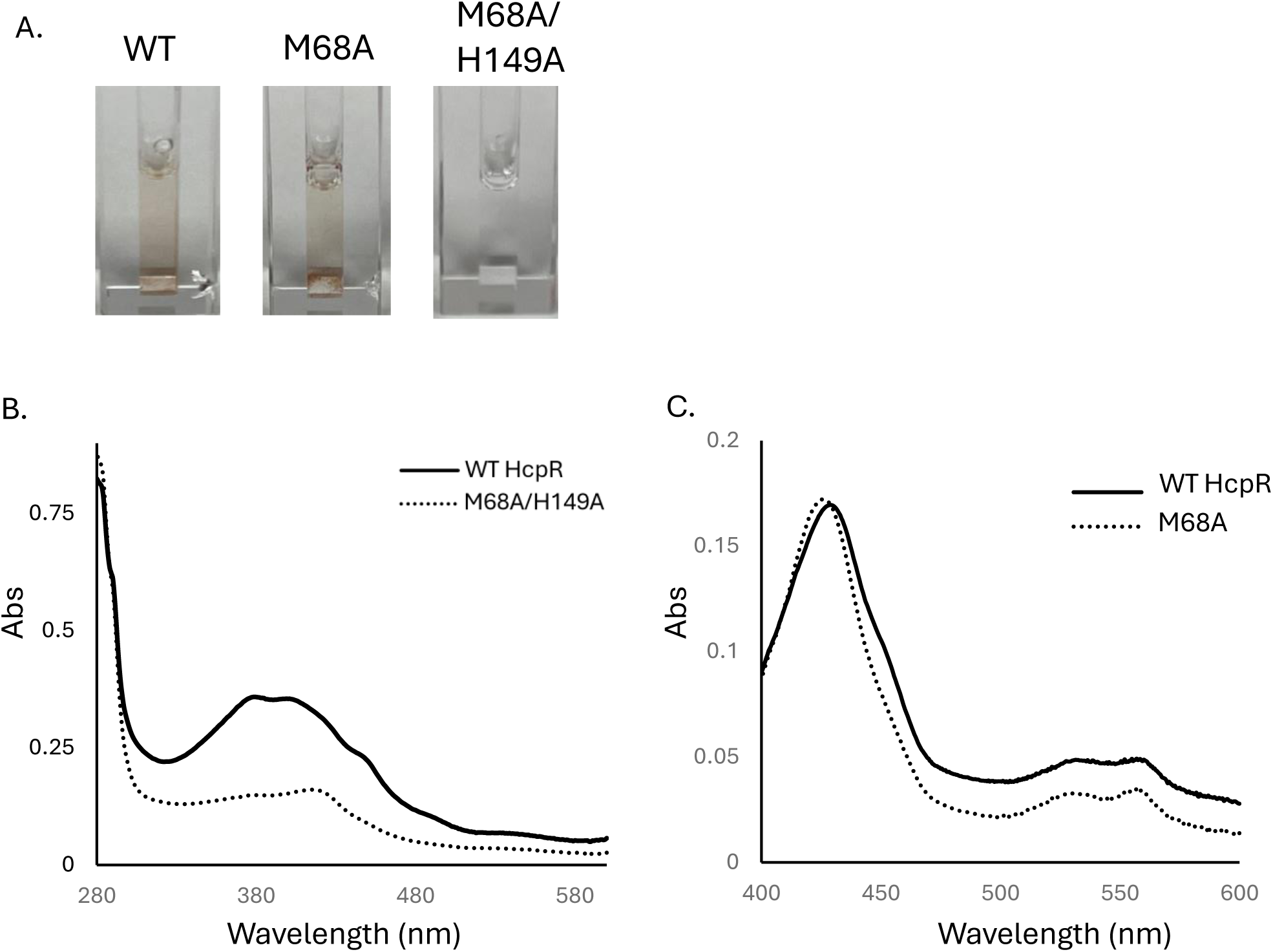
Met68 and His149 are necessary for proper binding of heme to HcpR. (A) Heme binding of WT, M68A, and M68A/H149A HcpR – the amber color of WT and M68A purified protein indicates the presence of heme in the sample. (B) Spectra of the reduced, heme bound WT-HcpR and the M68A/H149A double mutant HcpR. The loss of the Soret peak in the M68A/H149A HcpR indicates loss of heme binding. (C) Spectra of the reconstituted WT HcpR and M68A HcpR under reduced, anaerobic conditions. Changes in the location and form of the Soret peak indicates changes in the heme binding nature of the M68A HcpR.

To determine the importance of Met68 or His149 for HcpR activation, we mutated these residues to alanine and plasmid complemented an HcpR deficient strain of *P. gingivalis* (V2807) ^8^. Crucially, *P. gingivalis* lacking HcpR does not grow in physiologically relevant concentrations of NO_2_^−^, thus a strain with an inactive or deficient HcpR cannot tolerate elevated NO_2_^−^ levels. With treatment with NO_2_^−^ we saw a significant decrease in the growth of the M68A strain at 1mM NO_2_^−^ and no growth at the higher concentration of 2mM (Fig 4A). The double mutant of M68A/H149A could not grow at either concentration of NO_2_^−^ and was comparable to the ΔCHcpR negative control that completely lacks the DNA binding domain. Unexpectedly, with the H149A mutant we saw no significant decrease in growth at either concentration of NO_2_^−^.

To measure activity of the HcpR mutants, we assessed transcription of the *hcp* gene *in vivo.* In agreement with the growth studies, we saw a significant decrease of *hcp* transcript in the M68A complemented strain (Fig. 4B). However, there was still modest expression of *hcp*, which explains why the M68A strain exhibited some growth when compared to the L156* negative control at the lower concentration of nitrite. With the M68A/H149A double mutant we saw *hcp* transcript levels comparable to that of the negative control. With the H149A mutant we saw no significant decrease in *hcp* transcript levels.

To confirm expression and growth data we expressed and purified the M68A and M68A/H149A mutant HcpRs and compared them to wildtype HcpR spectroscopically. Both strains were purified at similar concentrations and purities to the wild type protein using cobalt affinity resin. The M68A/H149A double mutant did not bind heme effectively, which can be easily seen in the loss of color of the protein after reconstitution with heme and desalting (Fig. 5A). The UV-Vis spectrum revealed very little of the heme remained bound to the protein as evidence by lack of a Soret band (Fig 5B). The M68A mutant was still capable of binding heme and retains its coloring. However, there were subtle changes in the UV-Vis spectrum under anaerobic reduced conditions: there was a shift in the main Soret peak from 428nm in the wildtype HcpR to 426nm in M68A HcpR (Fig. 5C). There is also significant flattening of the α-β peaks at 533nm and 550nm. As the Soret spectrum arises from the heme-protein interactions, this indicates that there are changes in the heme binding pocket of M68A HcpR.

### Interdomain interactions are critical for generating a heme binding pocket

As we have shown, the formation of the heme binding site depends on the chain A DBD being in a conformation that allows it to form contacts with the flap region of Chain B. Arg226 of chain A is of particular interest, as it forms polydentate interactions with two backbone residues in the flap loop of chain B. The effect of this interaction is evident when comparing the FL-HcpR with the ΔCHcpR. The two structures superimpose with an RMSD of 0.564 Å (Fig. 6A). There is an ∼13.7 Å difference in the location of the flap region along residues 69-75 (Fig. 6B). These contacts are reflected in the B-factors of these regions in the two structures: the flap region of chain B in the FL-HcpR has lower B-factors than their corresponding regions in chain A (Fig. S5). The interdomain interactions create a divergence in the putative heme binding pocket size between the HcpR molecules in the dimer (Fig 6C). The pocket of FL-HcpR chain A aligns near perfectly with both chains of ΔCHcpR and is significantly smaller because there are no interactions between domains (Fig 6A). In both chains of ΔCHcpR and chain A of FL-HcpR, the pocket size is approximately 520 Å^3^.

**Figure 6.**
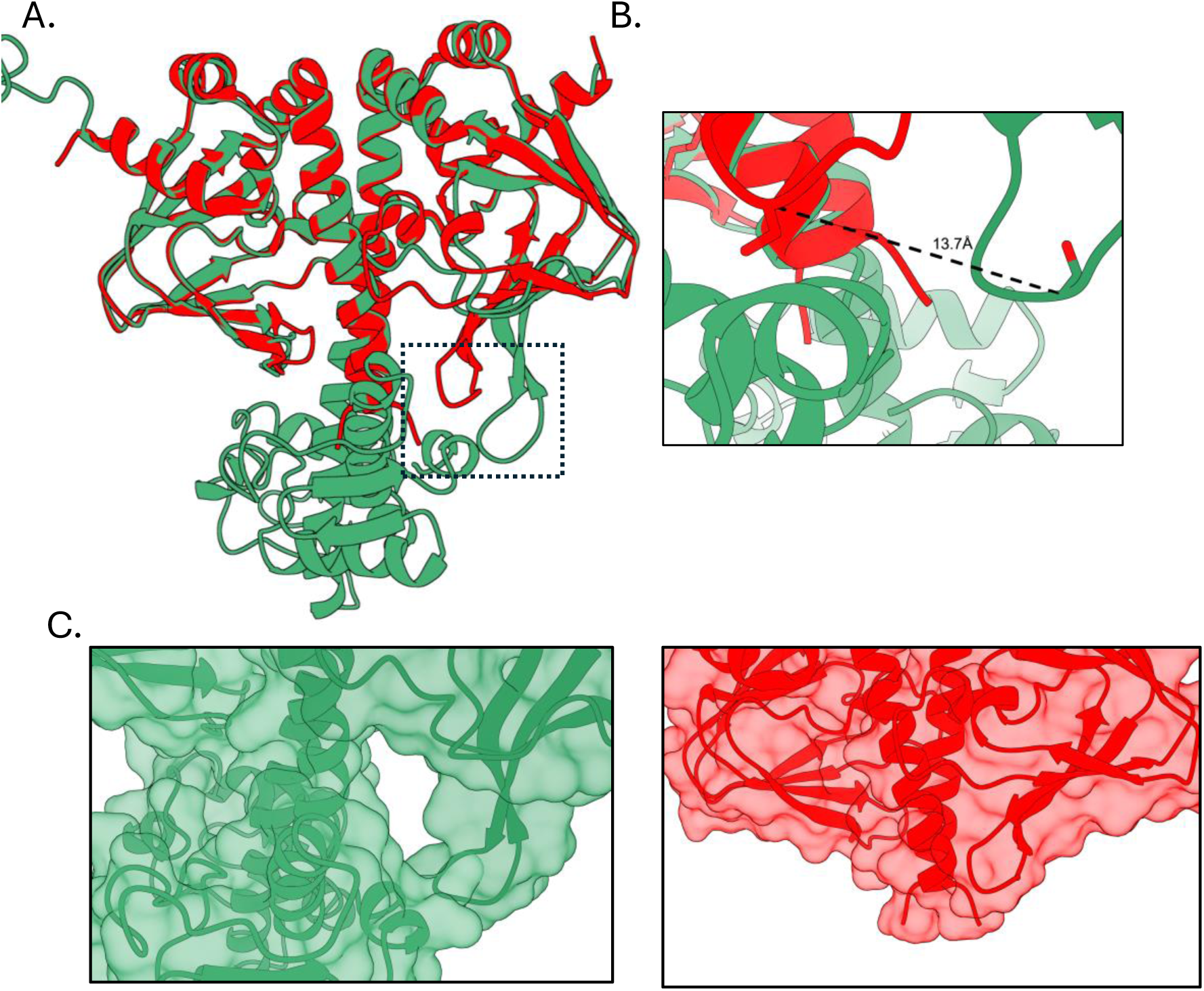
Structural alignment of full length HcpR and ΔCHcpR highlights the interdomain interactions that stabilize the heme pocket. (A) Structural overlay of the full length HcpR (green) and the C-terminal truncated ΔCHcpR (red). The structures overlay with an RMSD of 0.564Å. (B) Zoomed in view of boxed region in (A) of the flap region in chain B of HcpR compared to that of chain B in ΔCHcpR. The flap region of HcpR has shifted out by 13.7Å, enlarging the hydrophobic pocket. (C) Surface view of the boxed region in (A) of the pocket present in the full length HcpR and the subsequent position and ΔCHcpR.

In chain B of FL-HcpR, the pocket size increases to approximately 1100 Å^3^. This increased pocket size is much easier to accommodate heme, which has a volume of ∼510 Å^3^. This interaction demonstrates that the orientation of the DNA-binding domain can influence folding of the sensing domain. The difference in conformation between the two DBDs’ reflects the hinge region’s flexibility along residues 152-156 around which the DBDs move. In the crystal structure of FL-HcpR, the orientation of the DNA binding domain is also influenced by crystal packing forces (Fig. S6). These forces can stabilize the E helix in an extended form or allow the SD to form contacts with the DBD. When the DBD is allowed to contact the SD, as seen in the case of chain A DBD and chain B SD, the pocket is stabilized. In the crystal structure, the pocket that forms is further occupied by a portion of the N-terminus of an adjacent homo-dimer in the crystal lattice. Crucially, the pocket appears smaller in the SD of chain A and the ΔCHcpR structure due to the absence of the DBD interactions.

## Discussion

In this study, we present the structure of the full-length HcpR. The protein crystallizes in a peculiar form, with significant asymmetry in the C-terminal DNA binding domain that is not uncommon in CRP family regulators ^3^. The off-state may be represented by the extended E helix in chain B. This structure is present in both chains of the NO sensor DNR ^26^. However, it is unlikely that HcpR will adopt this structure in the off-state. The residues forming the leucine zipper motif that stabilize the homodimer do not extend past the hinge region at Leu153. After this point, most residues forming the motif have polar, hydrophilic side chains. Previously, we have demonstrated, using SAXS ensemble optimization modeling, that the DNA binding domain of Apo-HcpR can adopt several different conformations in solution ^27^. Thus, it is possible that the off form is an ensemble of different combinations, and activation of HcpR stabilizes the DNA binding domain into an orientation with accessible recognition helices.

The two conformations of the DBD can also be seen in some crystal structures of CooA and CAP. In HcpR, as in CooA, the DNA binding domain of chain B is in the fully “off” conformation with the recognition helix facing in towards the protein ^3^. The recognition helix of chain A is partially accessible; however, it is not in the fully outward facing orientation as seen in the structure of CAP bound to DNA. The heterogeneity of the two chains reflects the flexibility of the hinge region around which the DNA binding domains move and is important for creating the putative heme binding pocket. Moreover, the presence of the two conformations suggests that they are in a dynamic equilibrium in solution.

Guided by docking and mutagenesis studies, we interrogated the enlarged pocket in chain B as a putative heme-binding location. We identified two important residues in heme binding and activation, Met68 and His149. Both side chains are extended into the pocket and are in prime position to coordinate heme. Loss of Met68 severely inhibits the activity of HcpR. The single amino acid mutation decreases the activity by <10%. However, there is still a modicum of activity as measured using qRT-PCR. This may explain how the M68A complemented strain can persist at the lower nitrite concentration. Nitrite is bacteriostatic at these levels. It is possible a small amount of HcpR activity overtime can produce enough *hcp* transcript to allow for diminished growth. Loss of His149 does not affect the activity of HcpR. However, the M68/H149 double mutant HcpR has no activity, like ΔCHcpR. The purified M68A/H149A HcpR does not bind heme, explaining the activity observed *in vivo*. The His149 may play a role in heme coordination and stability of co-factor binding but appears to not play an indispensable role in the allosteric network that governs HcpR activation.

In the CRP family, distinct regulators have evolved to sense a wide variety of biologically relevant molecules involved in respiration, stress response, and metabolism. These regulators are based on a common modular design: (1) ligand binding β-barrels attached to a (2) central α-helix leucine zipper pair connected to a (3) C-terminal HTH DNA binding domain by a (4) flexible hinge region ^24,28–32^. Ligand binding appears to work on each of the modular parts, influencing their interactions and stabilizing the DNA binding domain in an “On” conformation (one in which the recognition helices are oriented such that the major groove of DNA is accessible). Despite their similar structural orientation and folds, activation depends on the exact nature of the ligand-induced reorganization of the β-barrel sensing domain and its effect on the allosteric networks communicating between the ligand binding domain and the DNA binding domain. Thus, among the CRP family members, key residues have little conservation (such as those involved in ligand interaction or co-factor binding).

This is reflected in the crystal structures and mechanisms of other well studied CRP members. In CAP, cyclic-AMP forms contacts with residues near the hinge region, extending the dimerization helix by 3 turns and forming a coiled-coil motif that stabilizes the orientation of the DNA binding domain ^33^. In CooA, binding of CO to heme displaces an N-terminal proline residue that frees up this region to form bridging contacts between the sensing domain and the DNA binding domain ^34^. In the halogen sensor CprK, ligand binding leads to a re-orientation of the binding pocket, decreasing its volume, that ultimately affects the position of key residues on the “flap” region that stabilize the DNA binding domain through formation of new contacts ^28^.

Despite CooA being the most well characterized heme binding CRP protein, several differences between CooA and HcpR are evident in the crystal structures. Although HcpR does have a proline residue near the N-terminus, the two helices adjacent to the N-terminus are well ordered and are not in a position that is located near the putative heme binding site. As previously mentioned, the extended loop region of HcpR corresponds to the heme binding location in CooA. In CooA, this region is shortened to accommodate the heme and is evident in the sequence of overlays of HcpR and CooA (FigS2). Thus, the heme binding location of HcpR is different from CooA and is likely “below” it (closer towards the C-terminal domain). The CooA mechanism of activation (the “Velcro” model) involves CO dependent displacement of the N-terminal proline residue bound to heme allowing the N-terminus to form contacts with the DNA binding domain ^34^. Based on the structure and heme binding studies here, it is likely that mechanism of HcpR is distinct from CooA.

DNR is a CRP family protein found in *Pseudomonas* that utilizes heme to sense NO, much like HcpR ^35,36^. DNR shares the highest sequence identity of the characterized CRP members with HcpR at 20%. The mechanism of DNR activation, although not well understood, differs from that of CooA. A significant conformational change in the crystal structure of DNR would be required for it to bind DNA, but the changes differ from those of CooA. Note that the position of the sensing domain in DNR structure is rotated by 60 degrees compared to the inactive form of CooA ^26^. Mutagenesis and heme binding studies reveal significant ligand switching occurs to accommodate mutations to heme coordinating residues ^37^. As there is little overlap in the conservation of key residues between DNR and HcpR it is difficult to speculate how their mechanisms may be analogous.

One possible mechanism for HcpR mirrors the pocket rearrangement CprK undergoes upon ligand binding. NO binding to heme displaces Met68. This displacement rearranges the pocket, shifting the flap region out along residues 72-75 into a position to make contacts with the DNA binding domain. These contacts stabilize the DNA binding domain into a position with the recognition helices accessible to DNA. This putative mechanism is attractive for several reasons. The location of the flap is very close to Met68 and displacement of the side chain can easily influence the flap location. This is bolstered by the highly conserved flap sequence in 72-75, with side chains that are accessible (Fig. 3). Among the CRP structures solved in the active formation, all adopt a similar DNA binding domain conformation in the active form. This would allow for the Ser72, Lys74, and Gln75 to form hydrogen bonds or salt bridges with the DNA binding domain. Furthermore, we have shown that this region capable of shifting when comparing the ΔCHcpR and FL-HcpR structures. This region is extended out by ∼14Å in the FL-HcpR structure, indicating the flexibility of this region to allow wide movements (Fig. 8B).

This can be illustrated using AlphaFold 3 to model the DNA binding domains in the active DNA bound form onto the crystal structure of FL-HcpR with heme docked (Fig. S6A) ^21^. The conserved flap residues form an interface with conserved residues adjacent to the recognition helix (Fig. S6B). This includes potential hydrogen bonding between Asn192 and Ser72 and the backbone at Gly73 and a potential network of ionic or hydrogen bonding interactions between Lys 74 and Gln75 and Lys183, Glu184, and Asp187. Additionally, Phe189 appears positioned at the entrance to the heme pocket, lending potential for hydrophobic interactions to occur here. As can be seen in a sequence overlay, the residues on the flap and the “turn” of the helix-turn-helix motif are highly conserved between different species of *Porphyromonas* and *Prevotella,* indicating their potential importance in stabilizing the active form.

Further experimental support is needed to confirm this hypothesis. Despite our attempts, we were unable to obtain viable crystals of the heme-protein complex for this report. Further studies involving the dynamics of NO binding and its effects on the nature of the heme pocket and DNA binding will help clarify the molecular mechanism of HcpR and the allosteric interactions that govern its activation.

---

**Figure S1 - Sequence alignments of HcpR found in *Porphyromonas* and *Prevotella* species.** Conservations of residues is graded on a scale of 1 to 11, where 1 indicates no conservation at that site and 10 (+) near identical conservation and 11 (*) indicates a residue is completely conserved across all sequences.

**Figure S2 - Arg226 and the flap region hydrogen bond.** Arg226 forms to hydrogen bonds with the flap region along the backbone of Val69 and Gly70.

**Figure S3 - Sequence alignment of CooA and HcpR.** The sequence alignment highlights the extended loop region of HcpR that occupies the heme binding site in CooA. Green highlight represents β-sheet structures and teal highlights represents α-helices.

**Figure S4 - Comparison of overall fold and conformational differences between HcpR and CAP.**(A) Stereo view of HcpR (green) and CAP (cyan) dimers depicting the overall organization of each protein. The N-terminal domain of HcpR (extending from residue 1-127) is larger than that of CAP (extending from residues 1-110). (B) Superposition of the HcpR-CAP overlay at the location of cyclic-AMP binding site. The extended loop region of HcpR (89-93) partially occludes this region.

**Figure S5 - B-factors in structure of HcpR.** Structural representations of B-factors on a scale of 18 (green)-148 (yellow).

**Figure S6 - Crystal packing in the full-length structure of HcpR.** The individual homo-dimer represented in blue and green chains with crystal contacts with adjacent homodimers in the crystal lattice.

**Figure S7 - Model of the full length HcpR structure in the active confirmation.** (A) Overview of the HcpR homo-dimer in the active confirmation bound to DNA. (B) Conserved residues that form the interface between the flap along residues 72-75 and the DBD along residues 183-192 with the sequence alignments of each region.

## Data Availability

The crystal structure data have been deposited to Protein Data Bank (PDB) with a title “Crystal Structure of the Transcriptional Regulator HcpR from *Porphyromonas gingivalis*” and assigned the following accession code(s): PDB ID 9DAY

## Funding information

This work was supported by NIH/NIDCR grant RO1DE023304 “Nitrosative Stress and Oral Bacteria (J. P. Lewis)”.

## Notes

### Competing Interest Statement

The authors have declared no competing interest.

### Summary of Updates

Minor adjustments to crystal structure space group and solution as well as the N-terminal sequence of the structure. This includes minor changes to Figure 1, Figure 2, and Figure 4.

