## Supplemental Figures for "Full-length structure and heme binding in the transcriptional regulator HcpR"

Fiigure S1

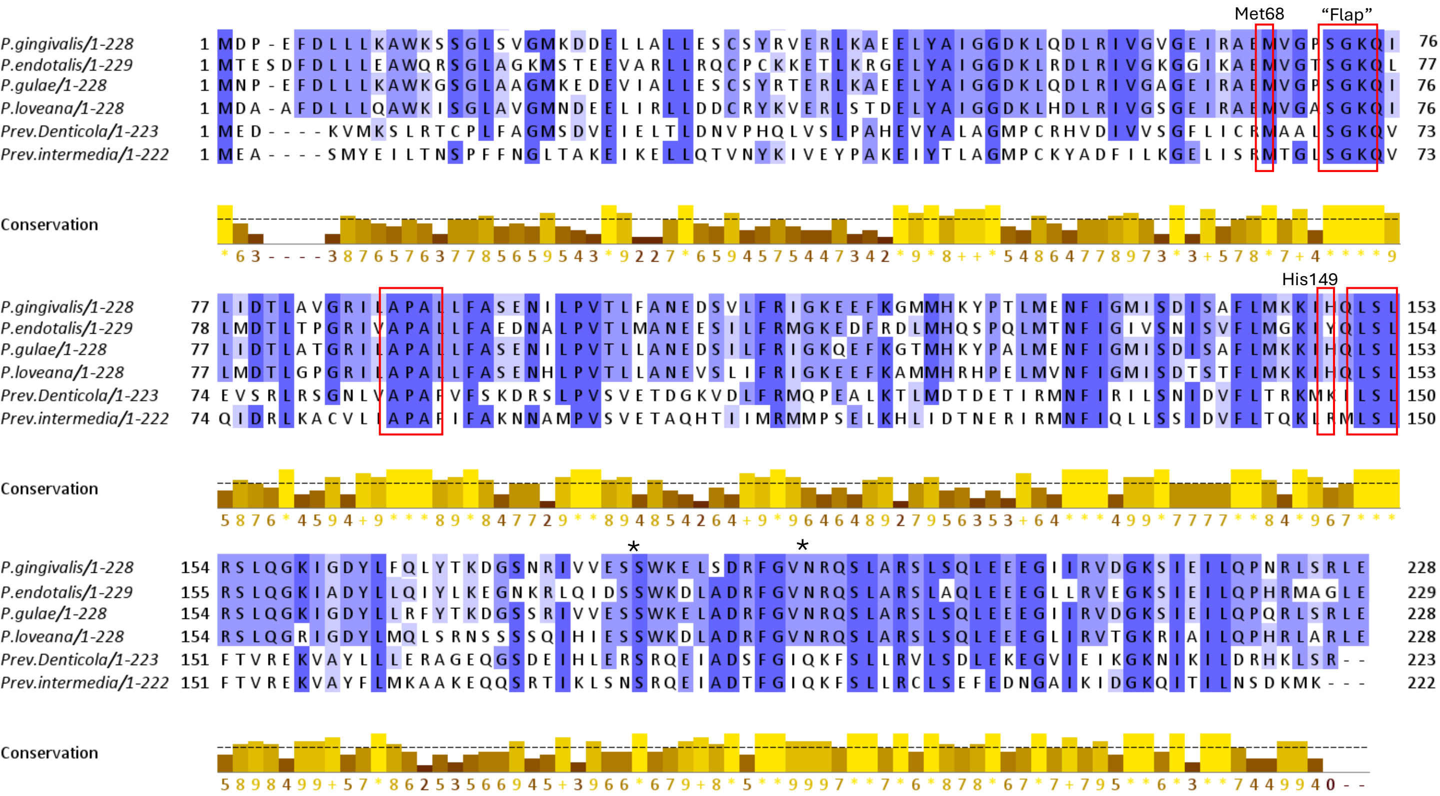

Met68
"Flap"
His149
P.gingivalis/1-228
P.endotalis/1-229
P.gulae/1-228
P.loveana/1-228
Prev.Denticola/1-223
Prev.intermedia/1-222
Conservation

Figure S2

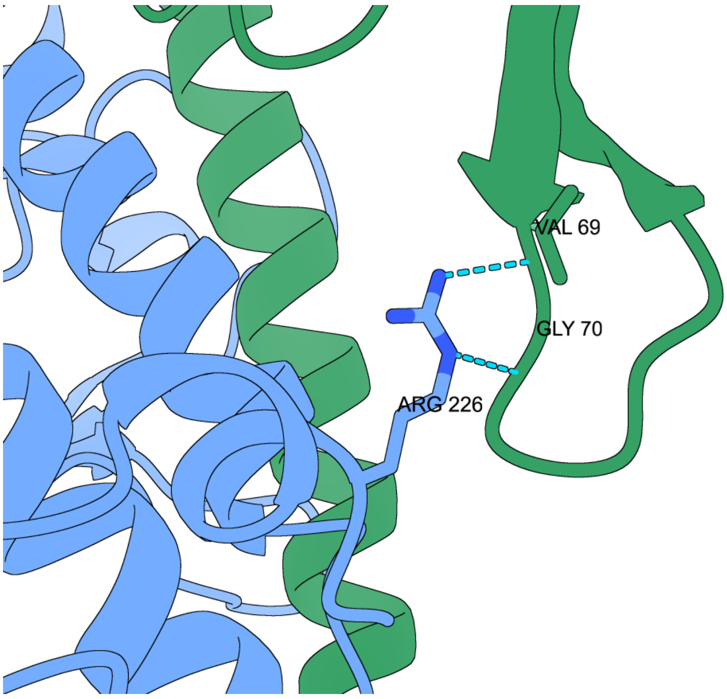

Figure S3

```
CooA        MPPRFNIANVLLSPDGETFF------------RGFRSKIHAKGSLVCTGE
HcpR        MDPEFDLLLKAWKSSGLSVGMKDDELLALLESCSYRVERLKAEELYAIGG
            * *.*::     ...* :.              .:* :     .* . *

CooA        GDENGVFVVVDGRLRVYLVGEE-REISLFYLTSGDMFCMHSGCLVEATER
HcpR        DKLQDLRIVGVGEIRAEMVGPSGKQILIDTLAVGRILA---PALLFASEN
            .. :.: :*  *.:*. :** . ::* :  *: * ::.    .*: *:*.
```

Loss of β6 in HcpR

```
CooA        -----------TEVRFADIRTFEQKLQTCPSMAWGLIAILGRALTSCMRT
HcpR        ILPVTLFANEDSVLFRIGKEEFKGMMHKYPTLMENFIGMISDISAFLMKK
                       : :   . . *:  ::. *::  .:*.::.   :  *:.
```

Loop extension in HcpR

```
CooA        IEDLMFHDIKQRIAGFFIDHANTTGRQTQGGVIVSVDFTVEEIANLIGSS
HcpR        IHQLSLRSLQGKIGDYLFQLYTKDG-----SNRIVVESSWKELSDRFGVN
            *.:* ::.:: :*..::::  .. *     .  : *: : :*::: :* .

CooA        RQTTSTALNSLIKEGYISRQGRGHYT-IPN-LVRLKAAADGDRDDDDD
HcpR        RQSLARSLSQLEEEGIIRVDGKSIEILQPNRLSRLE------------
            **: : :*..* :** *  :*:.     ** * **:
```

Figure S4

A.

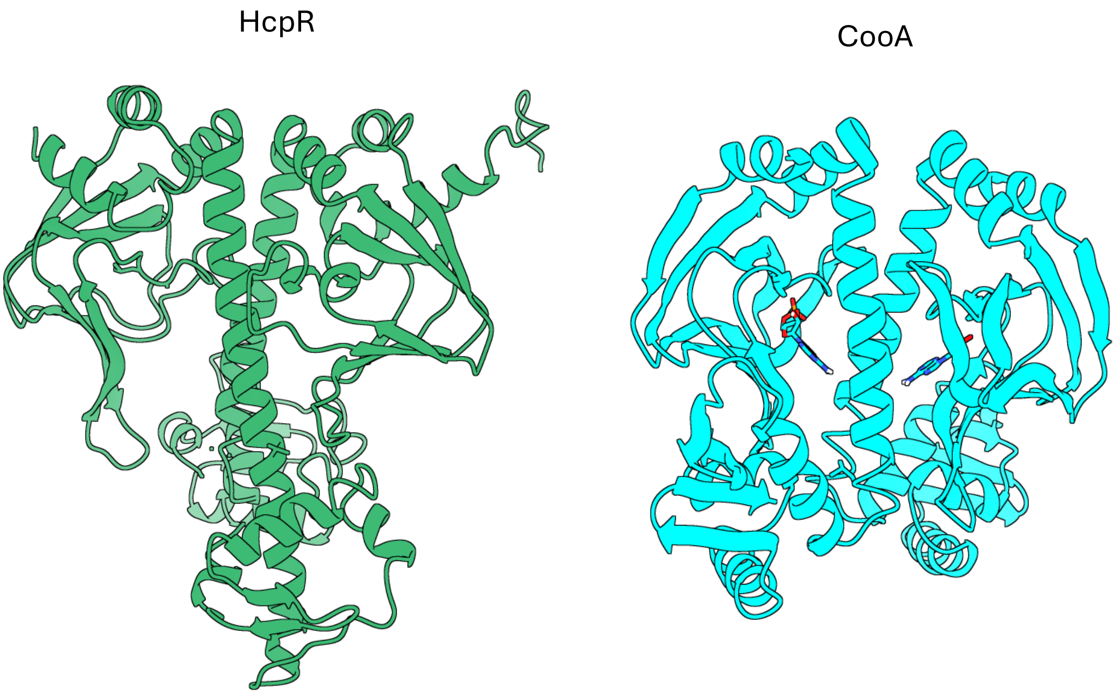


B.

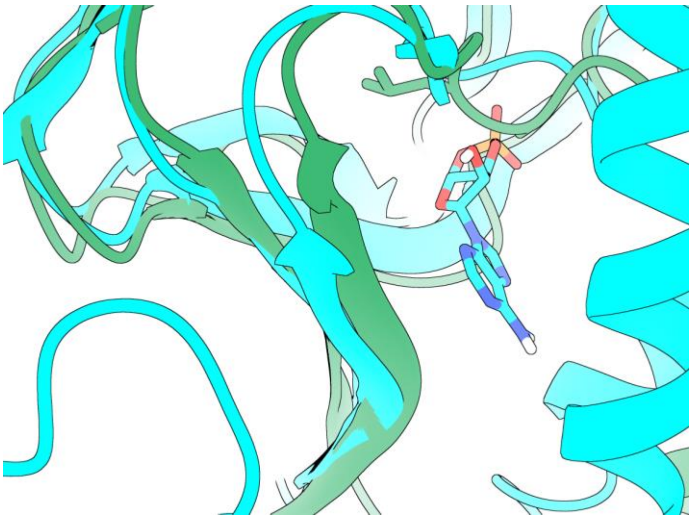

Figure S5

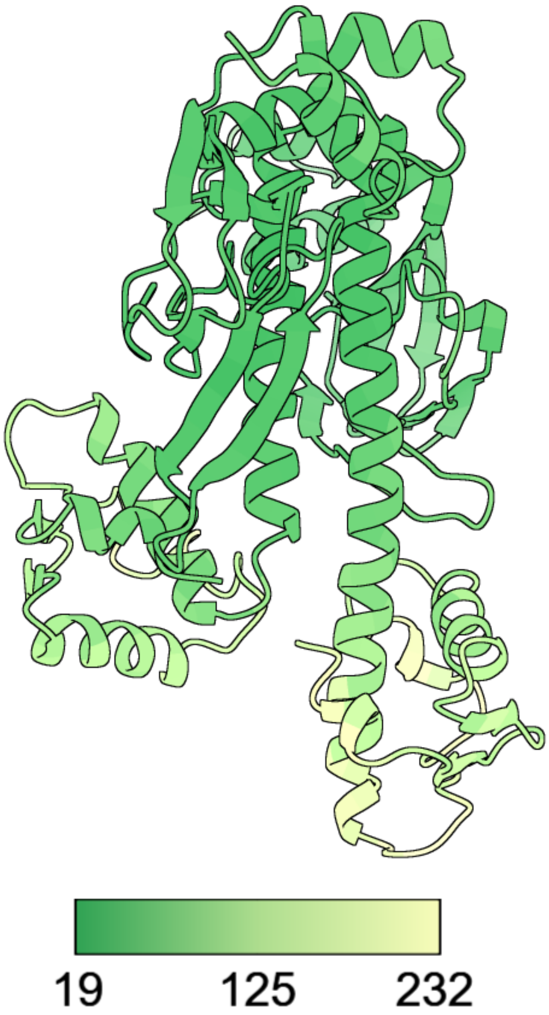

Figure S6

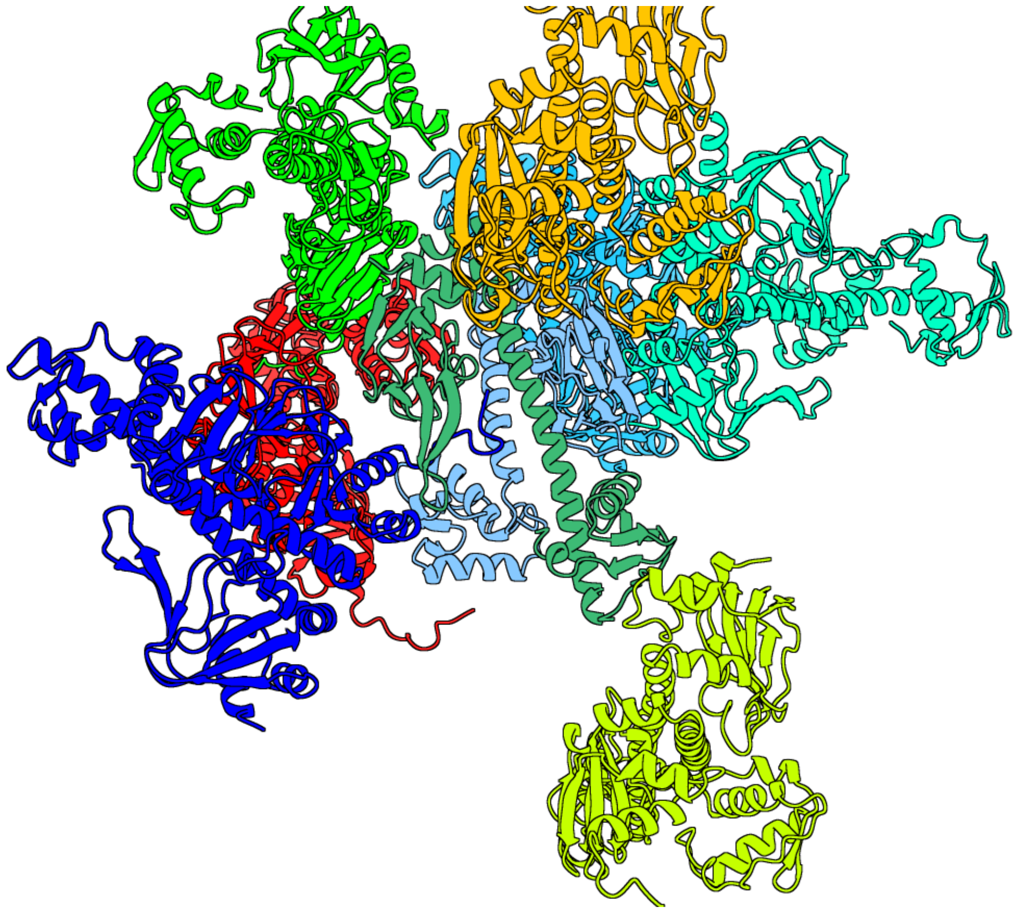

Figure S7

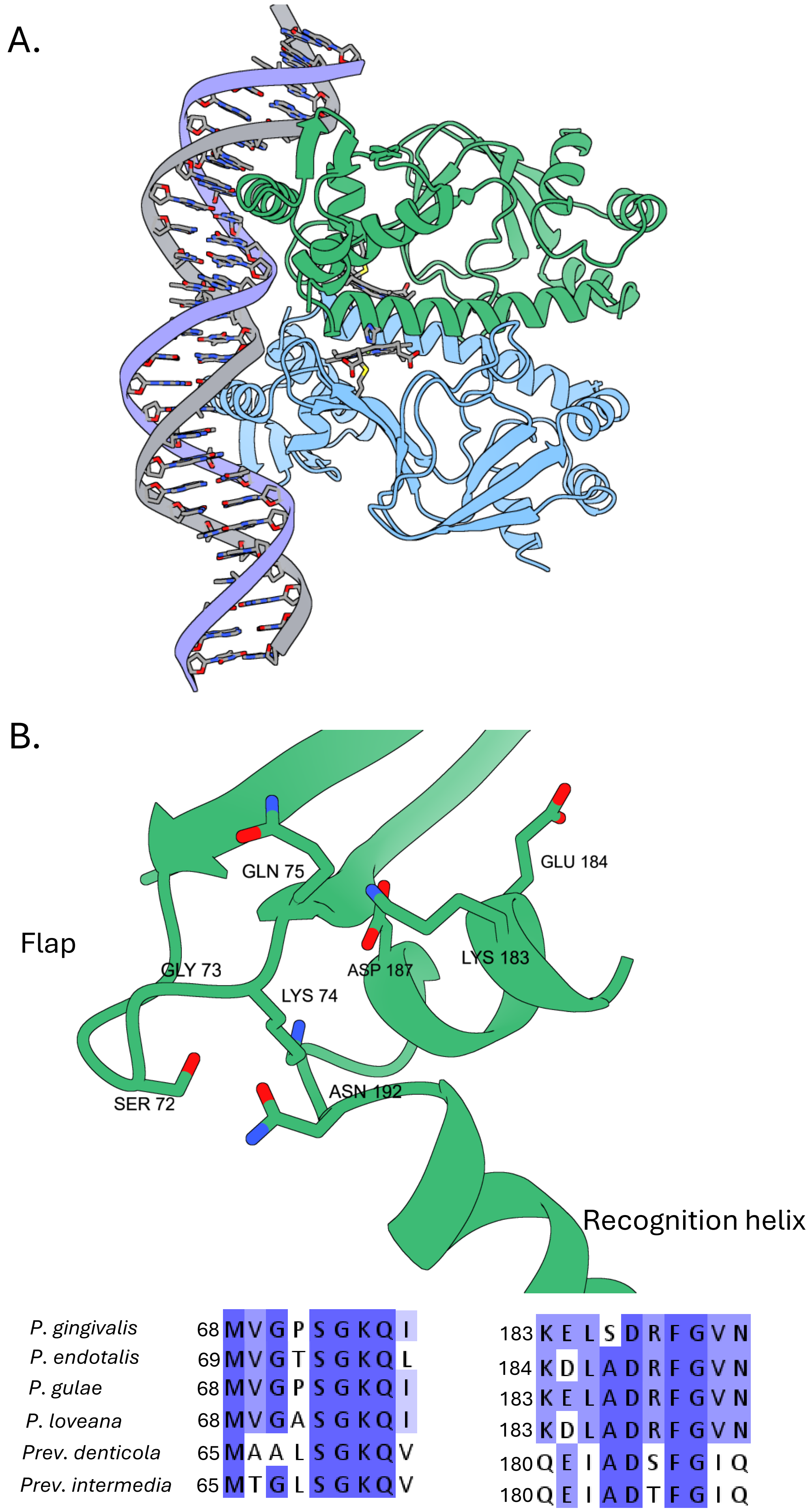
A.
B.
GLN 75
GLU 184
Flap
GLY 73
ASP 187
LYS 183
LYS 74
SER 72
ASN 192
Recognition helix
P. gingivalis 68 MVGPSGKQI 183 KELSDRFGVN
P. endotalis 69 MVGTSGKQL 184 KDLADRFGVN
P. gulae 68 MVGPSGKQI 183 KELADRFGVN
P. loveana 68 MVGASGKQI 183 KDLADRFGVN
Prev. denticola 65 MAALSGKQV 180 QEIADSFGIQ
Prev. intermedia 65 MTGLSGKQV 180 QEIADTFGIQ
